## Supplementary material for "Profiling of *Burkholderia pseudomallei* variants derived from Queensland’s clinical isolates": Supp Fig

Corresponding author:

**Supplementary Figures**

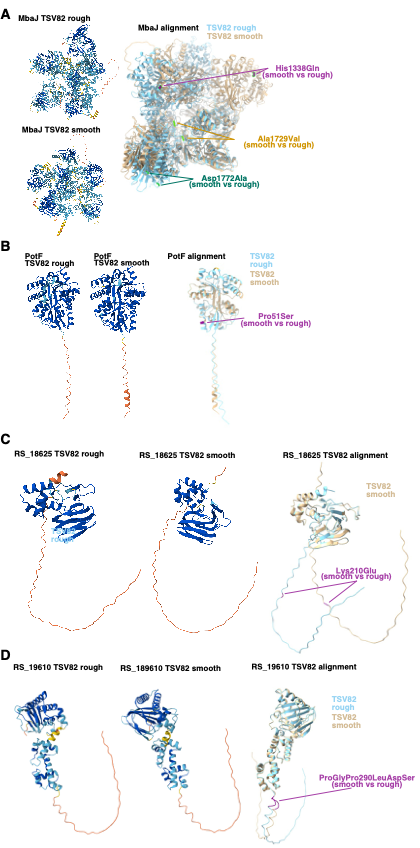

**Fig S1 Predicted structure of proteins carrying genomic mutations in TSV82 smooth compared to rough colony.** A) MbaJ B) PotF C) TSV82_RS_18625 D) TSV82_RS_19610.

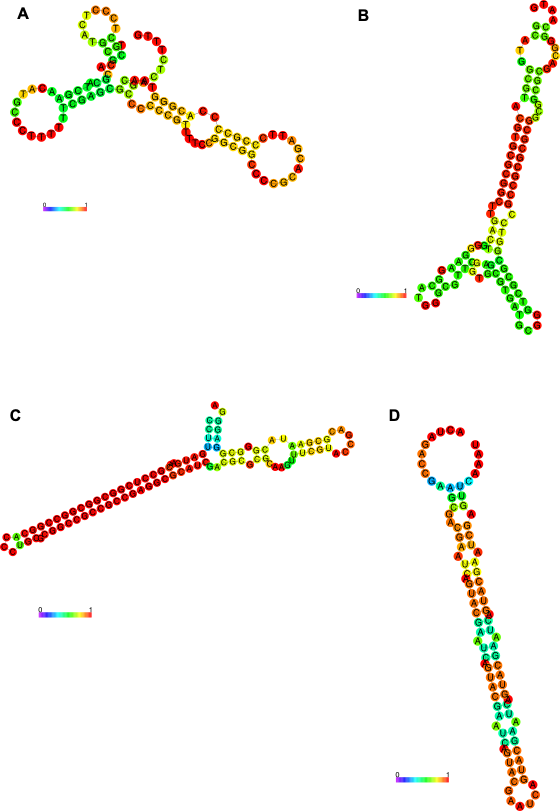

**Fig S2 Predicted secondary structures of candidate regulatory ncRNA found within regions of genomic variability.** A) Contig_1 2552774 Bp TSV82 B) Contig_2 538080 Bp TSV82 C) Contig_2 2200644 Bp TSV82 D) Contig_2 2170684 Bp TSV287. Scale bar represents the individual base-paired probability of the RNA transcript.
